# Synaptobrevin-2 C-terminal Flexible Region Regulates the Discharge of Catecholamine through the Fusion Pore

**DOI:** 10.1101/296582

**Authors:** Annita N. Weiss

**Author notes:** Address: The Max-Planck Institute for Biophysical Chemistry in Göttingen, Am Fa#x00DF;berg 11,37077 Göttingen.

## Abstract

The discharge of neurotransmitters from vesicles is a regulated process. Synaptobrevin-2 a SNARE protein, participates in this process through its interaction with other SNARE and associate proteins. Synaptobrevin-2 transmembrane domain is embedded into the vesicle lipid bilayer except for its last three residues. These residues are hydrophilic and constitute synaptobrevin-2 C-terminal flexible region. This region interacts with the intravesicular lipid bilayer phosphate head groups to initiate the fusion pore formation. Here it is shown that, this region also modulates the intravesicular membrane potential thereby the discharged of catecholamine. Synapotobrevin-2 Y113 residue was mutated to lysine or glutamate. The effects of these mutations on the exocytotic process in chromaffin cells were assessed using capacitance measurements, combined with amperometry and stimulation by flash photolysis of caged Ca^2+^. Both Y113E and Y113K mutations reduced the amplitudes of vesicle fusions and reduced the rates of release of catecholamine molecules in quanta release events. Further investigation revealed that the proximity of these charged residues near the vesicle lipid bilayer most likely changed the intravesicular potential, thereby slowing the flux of ions through the fusion pore, hence reducing the rate of catecholamine secretion. These results suggest that catecholamine efflux is couple with the intravesicular membrane potential.

## 1. Introduction

A series of successful events leads to the discharge of the vesicular content. These events include the priming of vesicles in which, vesicles that are docked at the cell plasma membrane await for the local influx of calcium to quickly fuse with the cell plasma membrane (1, 2). The priming step is followed by the formation of the fusion pore and the discharge of the vesicular content (3–5). Synaptobrevin/Vamp (sybII), a SNARE protein, participate in these events through its interaction with the vesicle lipid bilayer(6) and its interaction with other SNARE proteins, syntaxin and the synaptosome associated protein of 25 KD (SNAP-25) (7, 8). SybII N-terminal domain mutations inhibit the priming of vesicles and its C-terminal domain mutations reduce the speed at which vesicles discharge their contents (9, 10). SybII interaction with the vesicle lipid bilayer is accomplished through a set of mostly hydrophobic residues. This connection is increasingly been recognized as important for effective vesicle fusions. When sybII transmembrane region is partially deleted (11), is replaced by artificial lipid-anchors (12–14), or when it C-terminus is extended by polar residues (6), the fusion of vesicles is greatly inhibited or completely abrogated. Furthermore, the crystal structure of the SNARE complex that includes the transmembrane domains of sybII and syntaxin, shows that sybII and syntaxin form hydrogen bonds throughout their SNARE domains, linker regions and transmembrane domains. However, the last three residues of sybII C-terminus are devoid of such interaction and freely interacts with the vesicle lipid phosphate head groups (15). This region constitutes the sybII C-terminal flexible region. The relevance of this flexible region during the discharge of catecholamine in chromaffin cells was investigated.

## 1. Materials and Methods

### Single Cell Expression

Single chromaffin cells were isolated by enzymatic reaction from medulla glands of E17-E19 embryonic littermate. These double knockout (DKO) cells lacked both synaptobrevin-2 and cellubrevin. The cDNA encoding for sybII-Y113E and sybII-Y113K were produced by polymeric chain reaction (PCR) and verified by cDNA sequencing. Cells were infected using the Semliki Forest Viral expression system (16) 3 to 4 days after culture.

### Electrophysiology

Individual chromaffin cells were infected for 4 or 5 h to allow for the dual expression of eGFP and synaptobrevin-2. Flash photolysis and capacitance measurements were carried at whole-cell patch configuration by infusing 0.4 M fura-4F, 0.4 M mag-fura-2 (Molecular Probes, Eugene, OR) and 0.4 M CaCl_2_ bound to 0.5 M Nitrophenyl-EGTA. Ratiometric fluorescent measurements for the detection of calcium were performed as previously described (6).

Single vesicle capacitance measurements were performed at cell-attached patch configuration. The bath solution contained in mM: 145 NaCl, 1 MgCl2, 2.8 KCl, 2 CaCl2 and 10 HEPES, for which the pH and osmolarity were adjusted to 7.2 with NaOH and to 310 mOSM with D-glucose whenever necessary. The pipette solution contained in (mM) 50 NaCl, 100 TEA-Cl, 5 KCl, 5 CaCl2, 1 MgCl2, and 10 Hepes/NaOH (pH 7.2). The experiments were performed using the EPC-7 amplifier (HEKA) and the lock-in amplifier (SR 830; Stanford Research Systems). A 20-KHz at 50 mV sine wave stimulus at 100 mV/pA was used to resolve single capacitance steps. The phase was calibrated offline for each recording by using IgorPro software (Wavemetrics, Lake Oswego, OR).

### Amperometry

The high resolution amperometry measurement was used to detect the secretion of molecules from individual vesicles. The conventional carbon with a 10μM fiber diameter was used. To minimize errors due to diffusion, the carbon fiber was pressed onto the cell membrane and the tip of the electrode was cut before each data acquisition. The oxidation of molecules at surface of the electrode detected as amperometry spikes were acquired using EPC7, at the sampling frequency of 20 kHz and filtered at 3 kHz. To stimulate exocytosis, cells were infused at whole-cell configuration with a solution containing in (mM) 100 Cs-glutamate, 0.3 Na-GTP, 2 Mg-ATP, 2.5 CaCl_2_, 0.4 mM fura-4F, 0.4 mM furaptra, 20 DPTA and 32 HEPES, pH 7.2. The data are the mean of the median and One-way ANOVA was used for statistical analysis (*P<0.05, **P<0.01).

### Membrane Potential Calculation

The change in membrane intravesicular potential was calculated given that, prior to the merger of the vesicle membrane with the plasma membrane, the potential difference across the vesicle membrane (V_v_) and cell plasma membrane (V_c_) differs. However, when a connection between the two membranes is established, the vesicular membrane potential must reach the cell membrane potential resting state. The time dependence of this process was calculated according to:

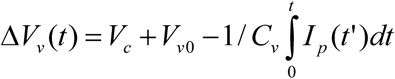

Where the cell membrane potential *V*_*c*_ =-70 mV, and the intravesicular potential prior to fusion *V*_*v0*_=+50 mV in the presence of ATP and Mg^2+^ (17). Therefore the initial condition for the intravesicular potential at t =0 is *V(t=0)=(V*_*c*_ + *V*_*v0*_). The vesicle capacitance change is modified by sybII C-terminal residues according to the energy that is required to move them from water to the membrane-water interface (6).

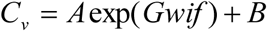

Where A=1 and B=0. The time constant for charging the capacitor was assumed to be 10&μs. The change in the intravesicular potential is facilitated by the flow of ions through the fusion pore(18). The current through the fusion pore due to the exchange of ion were calculated according the Nernst-Planck electrodiffusion equations using parameters as previously defined (19).

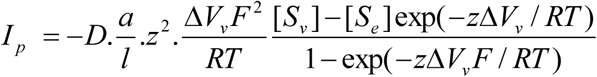

The concentration of catecholamine in a vesicle assume here was S_v_=135mM(19) and its external concentration S_e_=0mM

## 3. Results

In the full length SNARE complex, sybII is stabilized by its interaction with syntaxin. This is achieved through hydrogen bonds and Val der Waals surface interactions. However, its last three residues are polar and devoid of such interactions (grey, brown) [Fig. 1]. They form sybII transmembrane domain flexible region. In order to destabilize this flexible region, a single residue Y113, was mutated to lysine or glutamate. This location was chosen because prior investigation showed that charged amino acids that were added at the C-terminal end of sybII led to complete inhibitions of vesicle fusions (6). The gain-of-functions of the mutated constructs were assessed after their expressions in null double knockout (DKO) chromaffin cells. The fusion of vesicles with the cell plasma membrane was triggered by infusing photo-releasable calcium into cells at whole-cell patch configuration. The exocytotic response was monitored by parallel detection of the cell membrane capacitance and the release of the associated catecholamine by amperometry measurement.

**Figure. 1.**
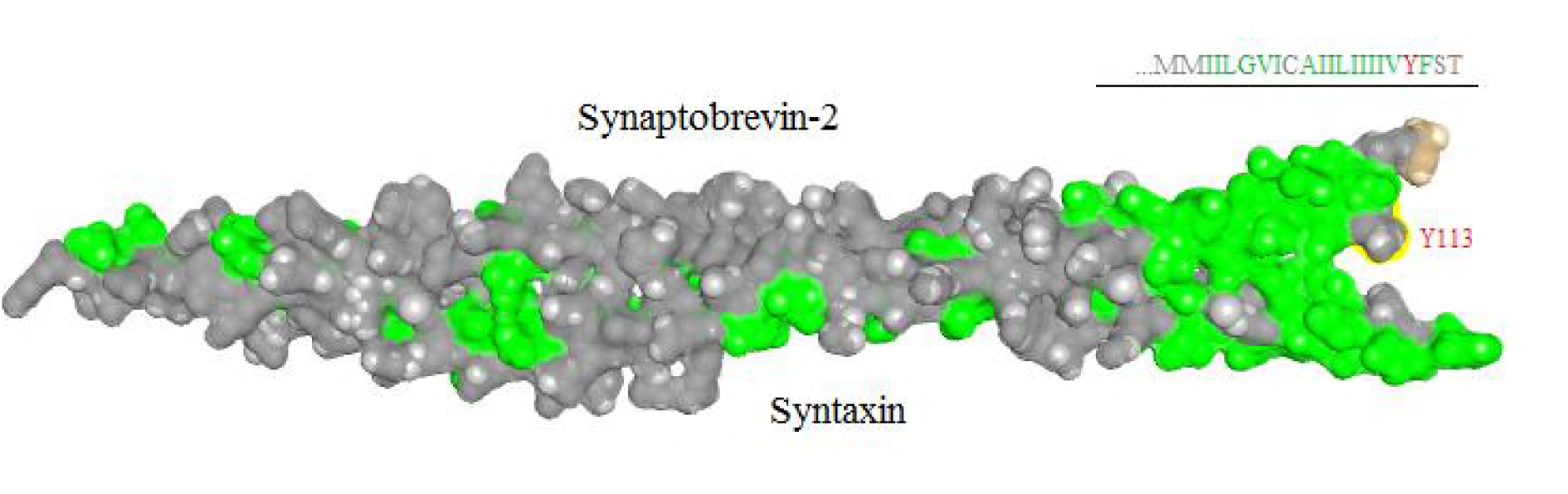
Van der Waals space-filling model of a helical bundle formed between syntaxin and sybil, including their transmembrane domains (PDB ID: 3IPD (15)). The polar residues are highlighted in grey, the residues in green are non-polar and sybil last residue threonine 116 is highlighted in brown. The hydrophilic region flexible sybil transmembrane region starts at residue Y113. This flexible region does not interact with syntaxin transmembrane domain after the SNARE complex is formed. Tyrosine residue at position 113 was mutated to single lysine (sybII-Y113K) and glutamate (sybII-Y113E).

### Charged residues in synaptobrevin-2 flexible region partially support vesicle fusion

SybII wild-type protein was expressed in DKO embryonic chromaffin cells. The exocytotic response to the rise of intracellular calcium was a biphasic increase of the capacitance trace. The capacitance trace could be sectioned into two components; the burst phase, representing the rapid change of the capacitance amplitude and the sustained phase its slower component lasting many seconds [Fig. 2A]. The burst and the sustained phases of cells expressing sybII-Y113K and sybII-Y113E were significantly reduced by ~50% of the capacitance amplitudes of cells expressing sybII [Fig. 2], suggesting an inhibition of vesicle fusion. Furthermore, when the capacitance traces were scaled to the same amplitudes at 0.5 sec after the calcium stimulus, only sybII-Y113E displayed a shower burst kinetic.

**Figure. 2.**
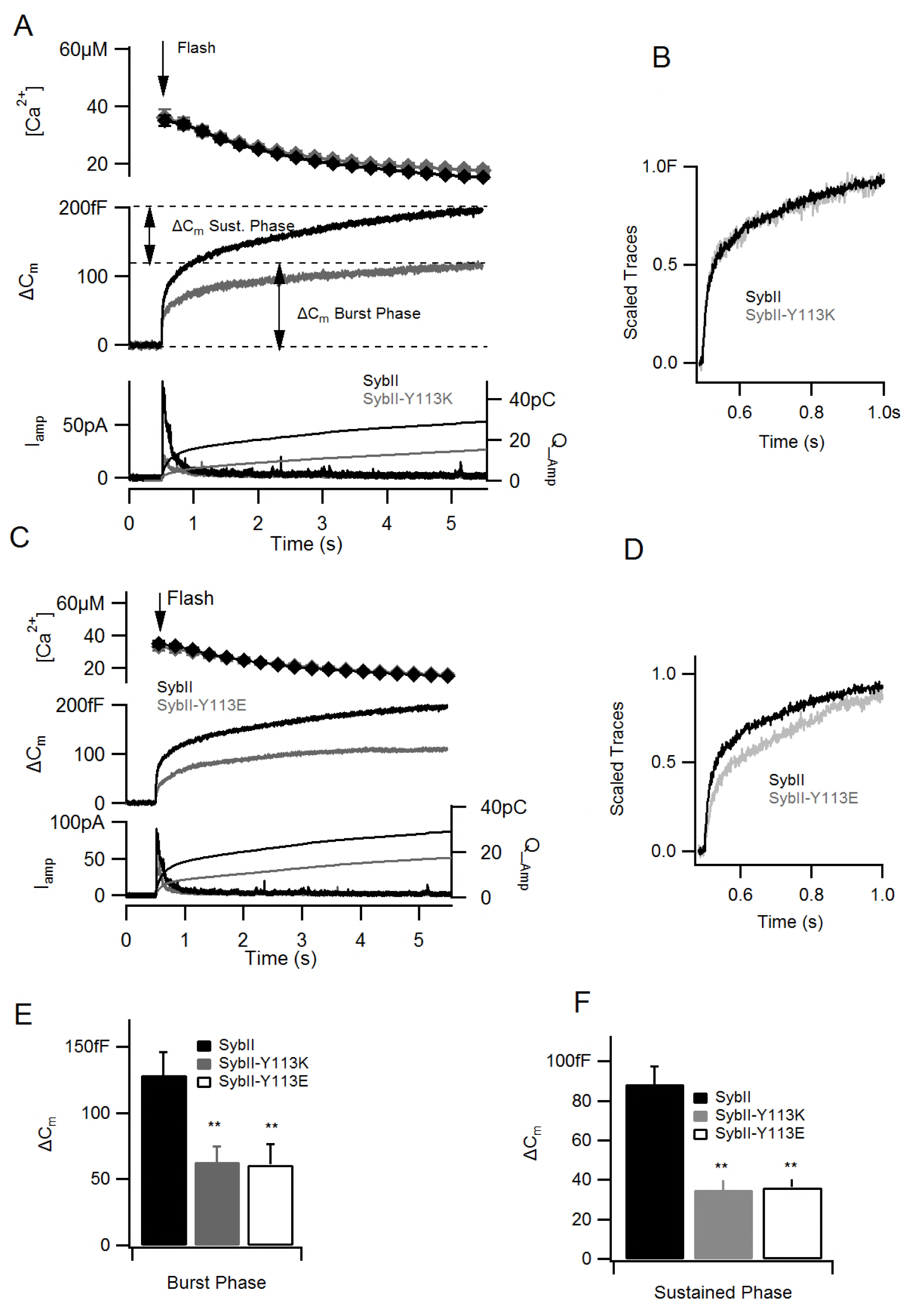
Charged residues within the transmembrane domain of sybil partially rescued the *exocytosis phenotype at whole-cell patch.* The fusion of vesicles was stimulated by releasing 2+ caged-calcium in patched cells. (A, C) At comparable [Ca^2+^]_i_ increase (top), cells expressing sybll-Y113K (n=20) and (B) sybII-Y113E (n=22) show a reduction of capacitance amplitudes (middle) and a reduction of amperometry currents (bottom) in comparison to sybll (n=18) overexpressing cells. (A) The capacitance amplitudes were biphasic. The burst phase was calculated as the capacitance change at 0.5 sec after the release of calcium (arrow). The sustained phase was calculated as the capacitance change between 0.5 and 5 sec. (E-F) Cells expressing Sybll-Y113K and sybll-Y 113E had a significant reduction of their burst and the sustained phases. (B-D) Only sybll-Y113E slowed the kinetic of the burst phase (*P<0.05, **P<0.01).

Next, individual vesicle sizes of cells harboring sybII-Y113K and sybII-Y113E were probed to determine whether the phenotype observed above could be explained by smaller vesicle sizes. The fusion of a single vesicle with the cell plasma membrane causes a stepwise increase of the capacitance when this one is probed at cell-attached patch clamp configuration. The current that was detected during these measurements were analyzed to extract the impedance properties (18). The real part (Re) of the patch admittance and the increase in the imaginary part (Im) were used to calculate the vesicle capacitance change (C_v_) [Fig. 3]. The mean±s.e.m capacitance step sizes of embryonic chromaffin cells expressing sybII (0.36 ±0.02fF) was similar to the average step size recorded in wild-type non infected cells (0.38±0.03fF). Also, cells expressing sybII-Y113K (0.34± 0.04fF) and sybII-Y113E (0.35± 0.03fF) had comparable step sizes as control. The average vesicle size of ~55nm calculated from these capacitance measurements is consistent with mouse embryonic vesicle sizes previously determined (20–22). This result indicates that neither mutation altered the size of vesicles.

**Figure. 3.**
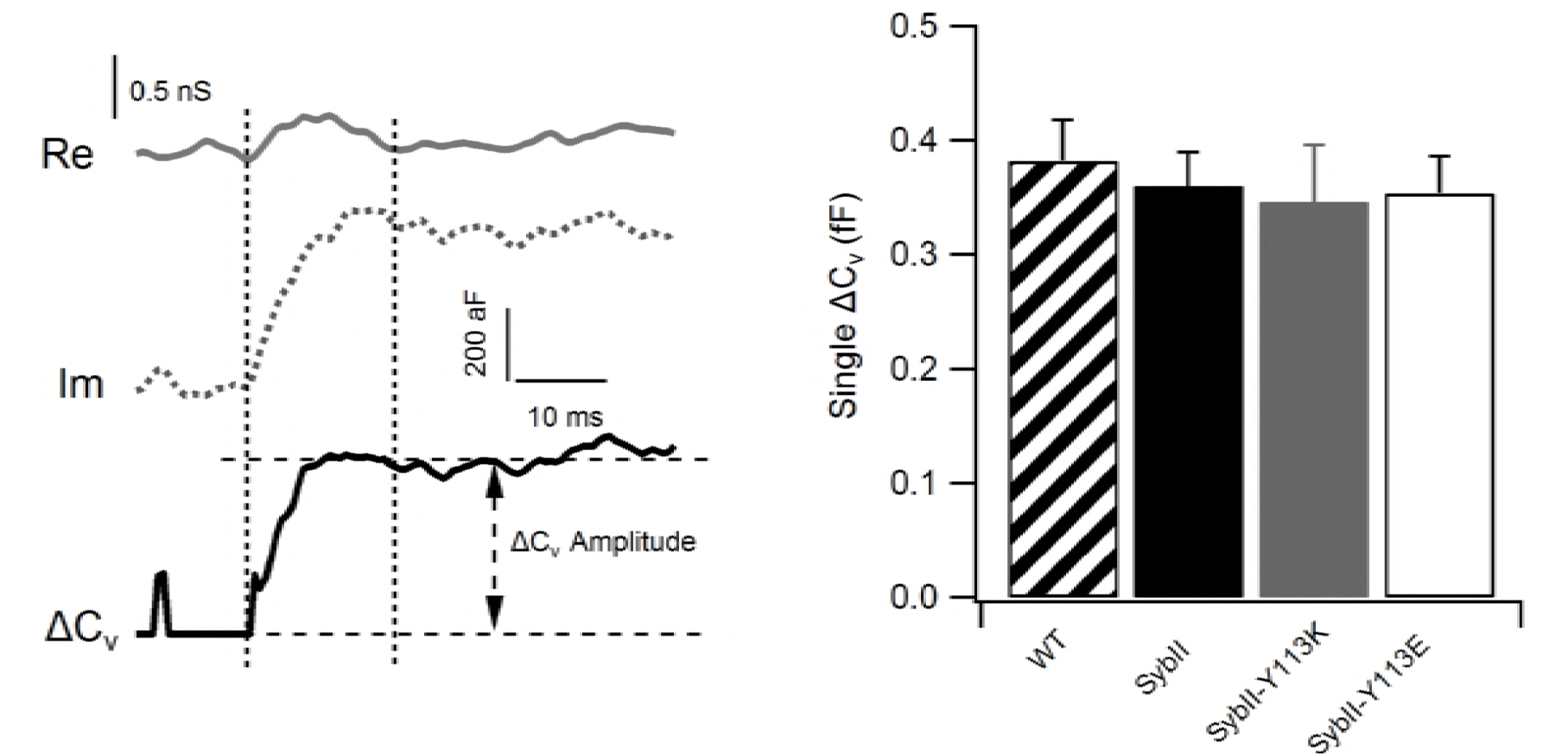
*Single capacitance measurement by cell-attached capacitance.* (A) Real part (Re, gray line) and Imaginary part (Im, dashed line) were calculated from the admittance measurement. The imaginary part and the real part provide information about the vesicle capacitance C_v_. Non-infected embryonic chromaffin cells (n=21 cells, 173 events) and cell expressing sybII (n=11 cells, 50 events), sybII-Y113K (n=14 cells, 52 events) and sybII-Y113E (n=11 cells, 79 events) all had similar vesicle capacitance step sizes.

### Synapobrevin-2 C terminus flexible region regulates the flux of catecholamine

An indirect mean to assess the fusion pore conductance is to monitor the flux of catecholamine during its discharged through the fusion pore. Therefore, it was investigated whether sybII C-terminal flexible region can modify the release characteristics of catecholamine. To detect the release of catecholamine molecules, a carbon fiber electrode was pressed onto the cell membrane. Catecholamine molecules secreted from vesicles oxidized at the surface of the potentiated carbon fiber electrode gave rise to amperometric current spikes [Fig.4B]. From these current spikes, the fusion pore duration and kinetics were extracted. More events/min were detected in cells expressing sybII (23 events/min) than in cells expressing sybII-Y113K (15 events/min) or cells expressing sybII-Y113E (17 events/min). The total amount of catecholamine molecules released from individual vesicles was similar regardless of whether the cells expressed sybII or the mutated proteins [Fig. 4A]. This is consistent with the above result which shows that these cells had similar vesicle sizes [Fig. 3] and also indicate that these mutations did not modify their contents. The mean current amplitude on the other hand differed. SybII-Y113E had the lowest peak amplitude (22.12 ± 2.40pA) followed by sybII-Y113K (36.99 ± 4.54pA), this is in comparison to (56.60 ± 6.19 pA) for sybII, indicating a slower diffusion of catecholamine molecules through the fusion pore in cells expressing the mutated constructs. Furthermore, sybII-Y113E had the broadest spike half-width [Fig. 4E], suggesting a delayed fusion pore expansion rate and an alteration of the fusion pore structure.

**Figure. 4.**
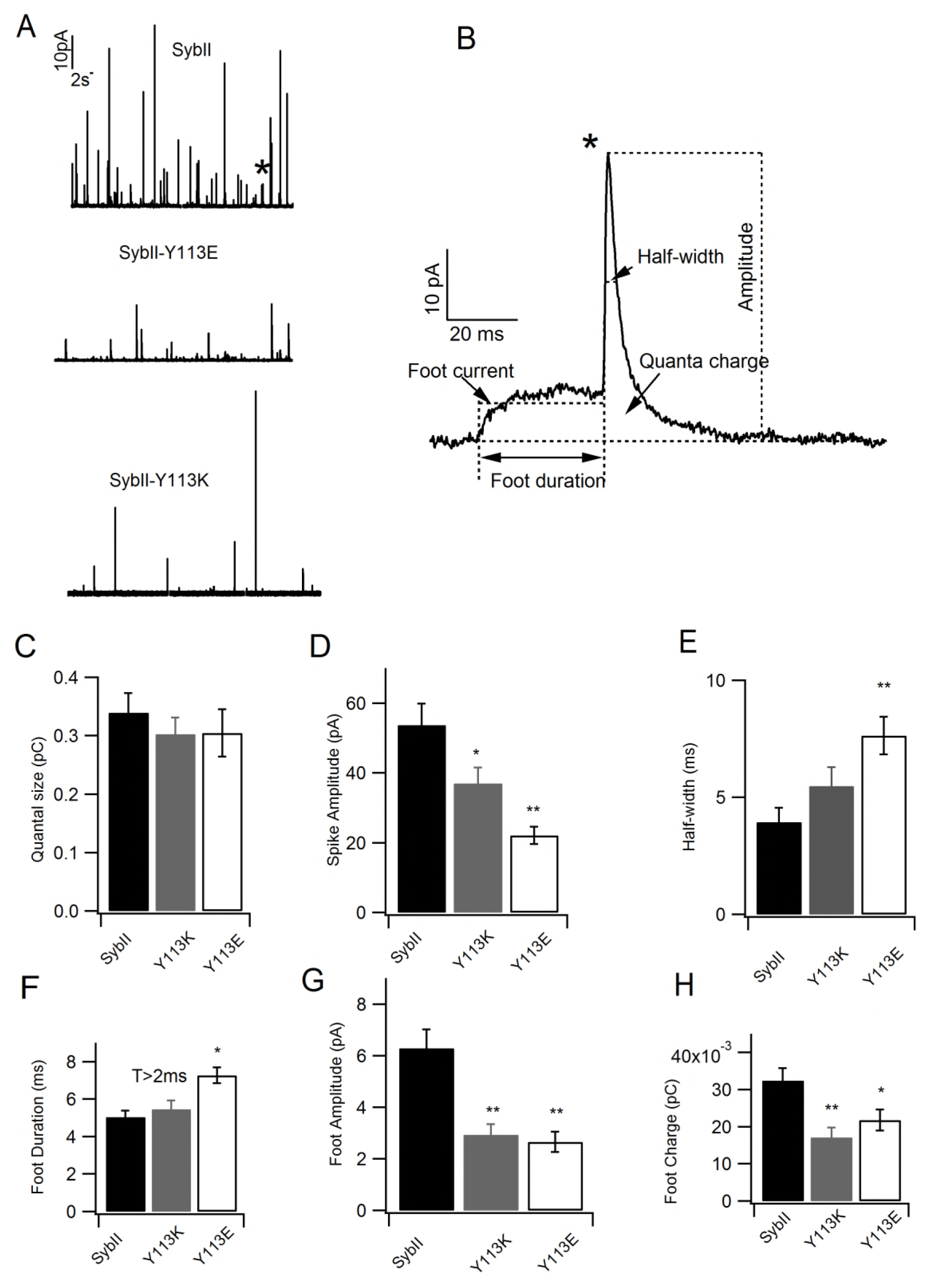
*SybII-Y113K and SybII-Y113E altered the fusion pore properties.* (A) Examples of amperometry traces recorded from cells expressing sybII (top), sybII-Y113E (middle) and sybII-Y113K (bottom). (B) Representation of a single amperometry spike with parameters as indicated (foot current, half-width, quanta charge, foot duration, and spike amplitude). C) Average quanta sizes are the same for all constructs. (D-E) Lower spike amplitudes and increase of spike half-widths in sybII-Y113E (n=17) and sybII-Y113K (n=17) in comparison to sybII (n=18). The foot parameters provide information about the state of the initial fusion pore. (F) Only the average foot duration of cells expressing sybII-Y113E was prolonged in comparison to that of sybII. (G-H) Whereas, the foot amplitudes and foot charges were reduced for both mutations (*P<0.05, **P<0.01).

The amperometry spike is often preceded by a & #x201C;foot” which reflects the formation of a metastable fusion pore that allows for the initial leakage of molecules. The foot characteristics reveal events that occur during the first milliseconds of the fusion pore expansion. The mean foot duration was longer for sybII-Y113K (7.27 ± 0.46 ms) and sybII-Y113E (7.27 ± 0.42 ms) than that of sybII (5.03 ± 0.34 ms), albeit only the foot duration of sybII-Y113E was significantly longer [Fig. 4F]. The foot amplitudes and the foot quanta release on the other hand were significantly smaller [Fig. 4G-H] for both mutations indicating that only few catecholamine molecules could diffuse through the initial fusion pore.

## 3. Discussion

The C-terminal end of sybII transmembrane region consists of hydrophilic residues. Unlike other domains of sybII, this region does not interact with syntaxin after the formation of the SNARE complex (15). This is likely to facilitate the interaction of sybII C-terminal region with the vesicle membrane lipid head groups and to facilitate the initiation of the fusion pore (6). In this study, tyrosine amino acid at position 113 of sybII transmembrane region was substituted to charged amino acids lysine and glutamate. These mutations still support the fusion of vesicles albeit to a lesser extent than that of the wild-type sybII protein, as depicted by a significant reduction of the burst and the sustained phases of the capacitance amplitudes. However, sybII-Y113K and sybII-Y113E did not modify the size of individual vesicles, indicating that fewer vesicles were fusion competent in these cells. The effects of sybII-Y113E and sybII-Y113K on the discharge of catecholamine were assessed by amperometry measurement. The foot amplitudes and foot charges were reduced for both mutations suggesting that the initial structure of the fusion pore was altered in both cases. The mutations also induced prolonged foot durations and spike half-widths albeit only those from sybII-Y113E expressing cells were significant, consistent with slower capacitance burst kinetic. These results indicate that fusion pore expansion and the discharged of catecholamine are slowed by these mutations and exclude any direct electrostatic effect of the side chains of lysine and glutamate on diffusing catecholamine molecules.

### Synaptobrevin-2 C-terminal flexible region can modify the intravesicular potential

It is observed here that, sybII-Y113E and sybII-Y113K reduction of catecholamine flux directly correlated with the nature of the substituted amino acids [Fig.5 A]. This is possible if these mutated residues altered the intravesicular potential, thereby reducing the flux of ions through the fusion pore, hence the secretion of catecholamine.

The amperometry measurement of the discharge of catecholamine from single vesicles suggests that the fusion of a vesicle starts with the formation of the fusion pore followed by its expansion. However, what is often overlooked is the fact that during this process the vesicle membrane which potential is initially distinct from that of the cell membrane potential must reach the resting cell membrane potential (18). Given the position of tyrosine 113 with respect of the lipid bilayer, its preferred location is the vesicle lipid bilayer region that contains water and lipid polar head groups (23, 24). At this position, sybII transmembrane domain residues would alter the intravesicular potential as they are incorporated into the membrane-water interface during the fusion pore formation as previously suggested (6).

The effects of lysine and glutamate side chains on the intravesicular potential during the initial fusion pore opening was assessed by calculating the intravesicular potential given that the capacitance change resulting from the vesicle fusion with the cell plasma membrane depends exponentially on the free energy of transfer (ΔGwif) from water to the water-membrane interface of sybII C-terminal residues (6) (Method). The free energy of transfer (ΔGwif) of tyrosine was used as the reference in these calculations. Since the energy of transfer (ΔGwif) of lysine and glutamate are higher than that of tyrosine (25), the intravesicular potentials initially at 50 mV (17) takes longer to equilibrate with that of the cell membrane potential (−70mV) for both residues [Fig. 5C]. This is due to the extra charges that are incorporated in the vesicle membrane. Furthermore, the intravesicular equipotential is also facilitated by the movement of ions through the fusion pore (18, 26), therefore, the flux of molecules through the fusion pore were calculated using the Nernst-Planck equation. This current was slower for lysine and glutamate [Fig. 5 C-B], suggesting that sybII mutations most likely slowed the equilibrium rate of the intravesicular potential, thereby also slowing the flux of ions through the fusion pore, hence the release of catecholamine molecules. This is consistent with previous findings that show that the extracellular reduction of sodium ions leads to reduced catecholamine flux through the fusion pore (19, 27). A similar effect was observed then membrane potential was increased in bovine chromaffin cells (28).

**Figure. 5.**
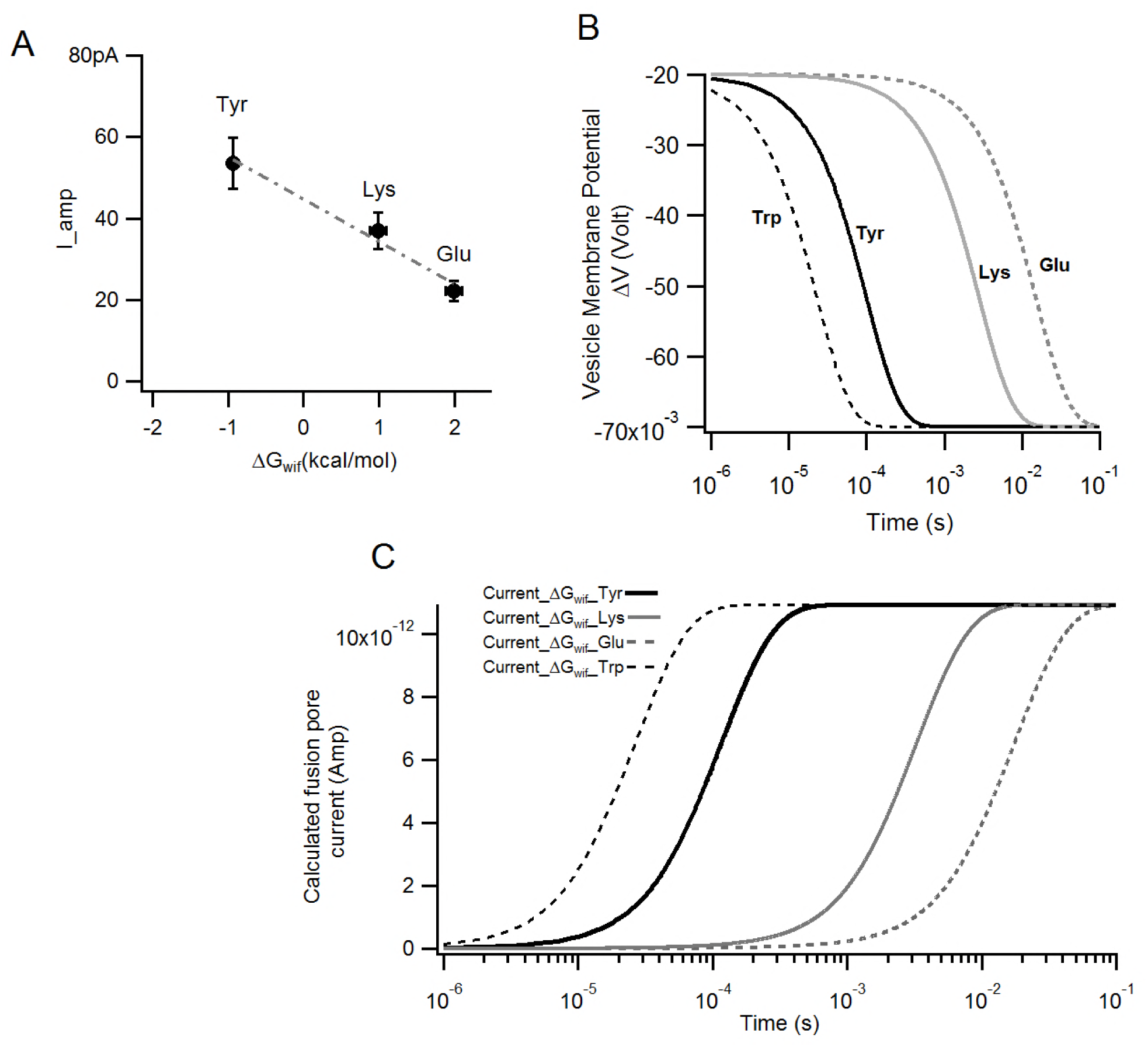
*Sybil transmembrane domain mutations sybII-Y113K and sybII-Y113E modify the intravesicular potential.* (A) The currents (I_amp) were experimentally determined (fig.4). They are the average spike amplitude for sybII, sybII-Y113K and sybII-Y113E. They are plotted against the energy that is required to transfer tyrosine, lysine or glutamate residues from water into the lipid bilayer interface (25). (B-C) The current and the changed in intravesiclular membrane potential were evaluated for residues tyrosine, glutamate, lysine and tryptophan, while taking into consideration the free energy of transfer of these residues from water into the water-lipid bilayer interface.Tryptophan were added for comparison since Y113 was previously mutated to tryptophan (ref, Jackson). These analyses indicate that as the fusion pore opens,the current flows through the fusion pore (B) and the intravesicular potential begins to change in order to reach the cell membrane potential. The intravesicular potential equilibrium is slowed when the energy of transfer (ΔG_wif_) of the mutated residues is higher than that of tyrosine.

In recent study, Y113 was mutated to tryptophan (21). Tryptophan is similar to tyrosine as it is found mainly at the water-lipid bilayer interface (24). Tryptophan in the membrane would speeds the intravesicular potential equilibrium [Fig. 5], thus speeding the fusion pore expansion, which is what was previously observed (21). The data presented here suggest that similarly to catechomlamine influx into the vesicle (17), its efflux is also couple with the intravesicular potential. Thus, the regulation of the voltage membrane potential is likely a physiological pathway to regulate calcium dependent exocytosis in chromaffin cells.

## Abbreviations

SybII: Synaptobrevin-2

## 4. Acknowledgements

I am grateful to Dirk Reuter and Ina Herfort for expert technical assistance and also grateful to Dr. Manfred Lindau and Dr. Jakob Sørensen for their critical reading of the manuscript. This work was supported by the National Institutes of Health grants R01NS38200, R01GM085808, T32GM007469, the Nanobiotechnology Center (a National Science Foundation Science and Technology Center, agreement No. ECS-9876771).

